# S3V2-IDEAS: a package for normalizing, denoising and integrating epigenomic datasets across different cell types

**DOI:** 10.1101/2020.09.08.287920

**Authors:** Guanjue Xiang, Belinda M. Giardine, Shaun Mahony, Yu Zhang, Ross C Hardison

## Abstract

**Summary:** Epigenetic modifications reflect key aspects of transcriptional regulation, and many epigenomic data sets have been generated under many biological contexts to provide insights into regulatory processes. However, the technical noise in epigenomic data sets and the many dimensions (features) examined make it challenging to effectively extract biologically meaningful inferences from these data sets. We developed a package that reduces noise while normalizing the epigenomic data by a novel normalization method, followed by integrative dimensional reduction by learning and assigning epigenetic states. This package, called S3V2-IDEAS, can be used to identify epigenetic states for multiple features, or identify signal intensity states and a master peak list across different cell types for a single feature. We illustrate the outputs and performance of S3V2-IDEAS using 137 epigenomics data sets from the VISION project that provides **V**al**I**dated **S**ystematic **I**ntegrati**ON** of epigenomic data in hematopoiesis.

**Availability and implementation:** S3V2-IDEAS pipeline is freely available as open source software released under an MIT license at: https://github.com/guanjue/S3V2_IDEAS_ESMP

**Supplementary information:** S3V2-IDEAS-bioinfo-supplementary-materials.pdf

## 1 Introduction

The tens of thousands of epigenomic datasets now available are potentially great resources to better understand the associations of epigenetic modifications with mechanisms of transcriptional regulation (ENCODE Project Consortium, 2012; Bernstein *et al*., 2010; Stunnenberg *et al*., 2016; Martens and Stunnenberg, 2013; Xiang, *et al*., 2020; Yue *et al*., 2014; Moore *et al*., 2020). However, integrating these resources for global inferences about regulation is challenging for many reasons. In this project, we focus on two issues. First, technical differences in procedures and biological samples analyzed in different laboratories introduce noise and biases that can obscure true biological differences (Shao *et al*., 2012; Xiang, *et al*., 2020; Meyer and Liu, 2014). Second, certain combinations of epigenetic modifications often appear together, but those combinations of modifications (epigenetic states) need to be learned from integrative modeling across epigenomic datasets simultaneously across multiple cell types (Ernst and Kellis, 2012; Zhang *et al*., 2016; Hoffman *et al*., 2012).

Here, we introduce a package, named S3V2-IDEAS, that builds upon our prior works and provides an improved, integrated workflow that will facilitate usability. In this pipeline, we address the first issue (noise and bias in data) by incorporating an improved version of the S3norm method (Xiang, *et al*., 2020), which can simultaneously normalize signals in foreground and signals in background. In contrast to S3norm, in which each 200bp bin was assessed as either foreground (peak) or background, in the improved version (S3V2) the reads within each bin are split into foreground reads and background reads. This strategy has been used in several previous studies (Mahony *et al*., 2014; Tarbell and Liu, 2019). After splitting reads, a single signal track can be converted into a foreground signal track and a background noise track. For the background noise track, both non-zero mean and non-zero standard deviations are matched across datasets, which can reduce the noise in some datasets (Fig. 1A, Supplementary Methods, and Supplementary Fig. 1D and 2). To address the second challenge (integration across multiple features and cell types), we performed genome segmentation using the Integrative and Discriminative Epigenome Annotation System (IDEAS), which learns epigenetic state models from the normalized epigenomic signals simultaneously along the genome and across cell types to improve consistency of state assignments across different cell types (Zhang *et al*., 2016); Zhang and Hardison, 2017). Moreover, the IDEAS model can jointly estimate the state of a genomic region by using the information in a set of similar cell types, so that the state can be accurately estimated even for cell types with missing data (Zhang and Mahony, 2019). The S3V2-IDEAS pipeline incorporates both S3V2 normalization and IDEAS segmentation so that the advantages of both methods can be used to normalize, denoise, and integrate multidimensional epigenomic datasets across different cell types.

**Figure 1.**
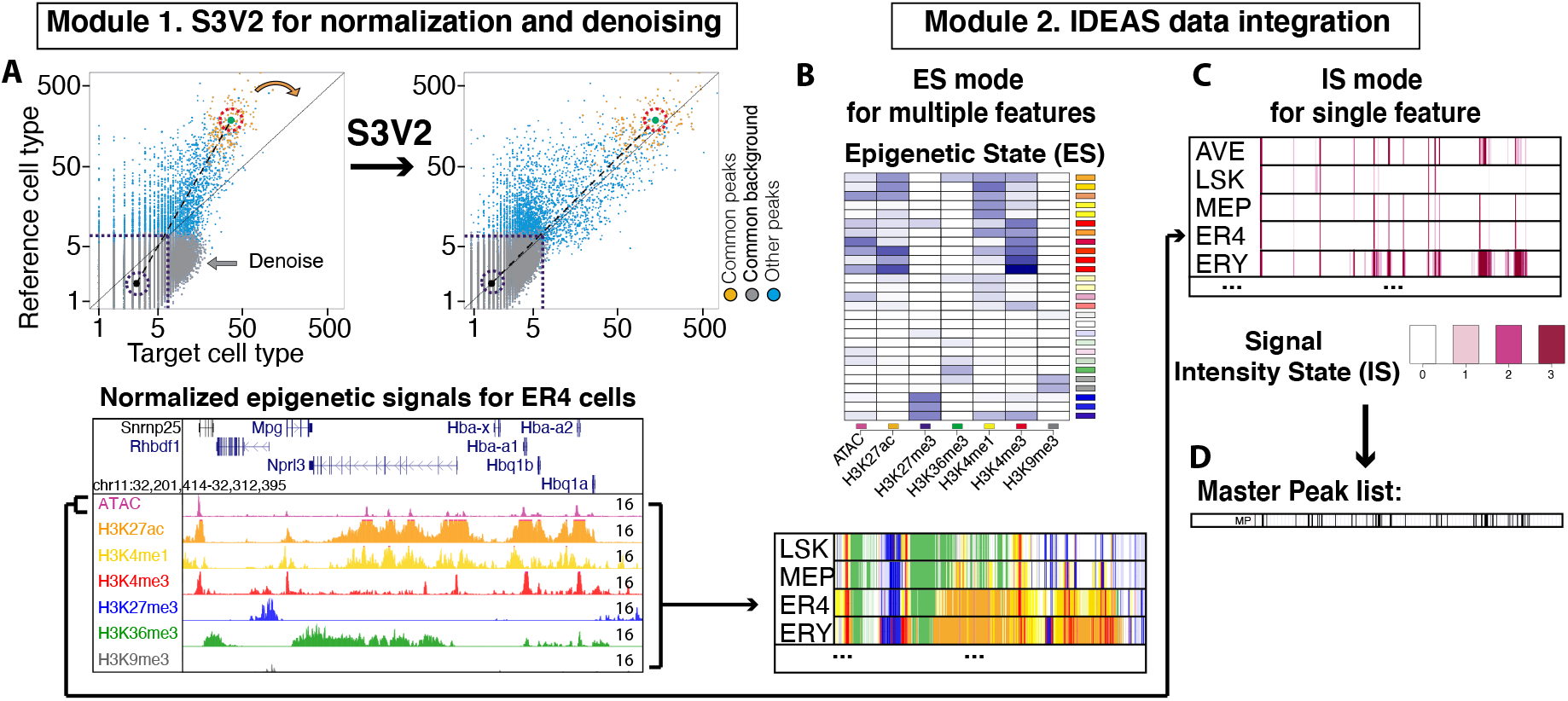
Overview of S3V2-IDEAS pipeline. (A) Module 1 normalizes and denoises input data using the S3V2 method. Examples of normal-ized epigenetic signals from the Hba locus in G1E-ER4 cells (ER4) are shown. (B and C) In Module 2, the normalized data is integrated by IDEAS in one of two modes. (B) The epigenetic state mode integrates multiple epigenetic features into an epigenetic states model. (C) The signal intensity state mode finds frequently occurring signal intensity states for a single epigenetic feature, along with a master peak list derived from those states (D). AVE = average, LSK, MEP, ER4, ERY = abbreviations for cell types (Xiang, Keller, Heuston, *et al*., 2020).

## 2 Implementations

The inputs to S3V2-IDEAS are (1) average read counts of each epigenetic feature in each cell type (bigWig), (2) an annotation file that includes the names of the cell types and the epigenetic features of bigwig files, and (3) information about the mapped genome, such as chromosome sizes and black-listed regions (Kent *et al*., 2002, 2010; Amemiya *et al*., 2019; Boyle *et al*., 2014; Yue *et al*., 2014).

The S3V2-IDEAS incorporates two major modules. First, it uses the S3V2 method to normalize and denoise the epigenomic datasets (Fig. 1A). The second module of the package incorporates the IDEAS genome segmentation model to integrate the epigenomic signal into tracks of epigenetic state assignments for each bin in each cell type (Fig. 1B and C). The second module can operate in either of two modes. When the input data include multiple epigenetic features, the module executes an epigenetic states mode (ES mode), which integrates the signals of multiple epigenomic features into epigenetic states as done previously (Fig. 1B). When the input data include one epigenetic feature, the module executes a signal intensity state mode (IS mode) to learn the most frequently occurring states of that one epigenomic feature (Fig. 1C). In the IS mode, a master peak list (Fig. 1D) can be extracted from the signal intensity state tracks by a novel hierarchical method which provide way to integrate the epigenomic information across cell types (Supplementary Figs. 5 and 6).

## 3 Results and Discussion

The S3V2-IDEAS produces three outputs: the normalized signal tracks and the −log10 p-value tracks based on the background model (Fig. 1A); a list of epigenetic states or signal intensity states and the corresponding state track in each cell type (Fig. 1B and 1C). An additional master peak list can be produced in the IS mode (Fig. 1D).

To illustrate these results, we applied the S3V2-IDEAS to datasets compiled by the ValIdated and Systematic integratION of epigenomic data project (VISION) (Xiang, *et al*., 2020; Heuston *et al*., 2018; Hardison *et al*., 2020). The ES mode can integrate seven epigenetic features to a 27 epigenetic states model (Fig. 1B). Compared with our previous analysis (Xiang et al. 2020), the genome segmentation tracks from this S3V2-IDEAS are more consistent between biological replicates (Supplementary Fig. 2C and D).

We illustrate the IS mode by limiting our analysis to only the ATAC-seq. In the IS mode, the ATAC-seq signal tracks can be first normalized and converted into tracks of signal intensity state (Fig. 1C). Then, a master peak list can be extracted from these state tracks (Fig. 1D). A master peak list is a straightforward way to obtain a coherent set of chromatin accessible peaks across cell types, which can be challenging for larger numbers of cell types (Meuleman *et al*., 2020). Comparing with the one produced by simply pooling and merging the MACS2 peaks in all cell type (Zhang *et al*., 2008), the IS mode master peak list pinpoints functional elements with higher accuracy (Supplementary Fig. 6 and 7).

These results indicate that the S3V2-IDEAS should be versatile and effective tool for integrative analyses of epigenomic datasets.

## 4 Funding

The work is supported by NIH grants GM121613 and DK106766.

## Supplementary Data

### 1. Overview of the S3V2-IDEAS package

S3V2-IDEAS is an easily applicable package that normalizes, denoises and integrates epigenomic data sets across different cell types. This package has two modules. The first module develop an improved version (S3V2) of our previous developed S3norm method to normalize the epigenomic data sets across different epigenetic features and different cell types (Xiang, Keller, Giardine, *et al*., 2020). Comparing with the S3norm, S3V2 can first split the signal track of each dataset into a foreground signal track and background noise track. For the background noise track, both non-zero mean and non-zero standard deviation can be equalized across different datasets, so that some datasets can be denoised. Within the first module, a dynamic negative binomial model is incorporated to adjust for variation of local background (Zhang *et al*., 2008; Xiang, Keller, Giardine, *et al*., 2020). The second module uses a two-dimensional IDEAS genome segmentation model to integrate the normalized epigenomic signal tracks into state tracks (Zhang *et al*., 2016). To increase the versatility of IDEAS, we incorporated two mode in the second module. When the data inputs include multiple epigenetic features, the module integrates those multi-dimensional signals into a list of epigenetic states (ES model), with each state representing a distinct combination of epigenetic features. When the input data consist of only one epigenetic feature, the module executes a signal intensity state mode (IS mode) to learn the frequently occurring states of signal strength. In both modes, a set of genome segmentation tracks is generated by assigning an epigenetic state or a signal intensity state to each genomic region in each cell type. To facilitate the downstream analysis, such as analyzing the epigenetic events across different cell types, we further developed a novel method to extract a master peak list from the signal intensity state tracks across all cell types. We describe each of these steps in more details in the following sections.

### 2. Data preprocessing

To illustrate the usage of the S3V2-IDEAS, we used the this package to analyze the epigenomic data sets compiled by the ValIdated and Systematic integratION of epigenomic data project (VISION: usevision.org), which includes seven epigenetic features (H3K4me3, H3K4me1, H3K27ac, H3K36me3, H3K27me3, H3K9me3 and chromatin sensitivity) in 20 hematopoietic cell types in mice (Xiang, Keller, Heuston, *et al*., 2020; Heuston *et al*., 2018; Hardison *et al*., 2020). The mapped reads in these data sets were processed by the data preprocessing pipeline in the VISION project. We then divided the mm10 mouse genome assembly into ~13 million 200-bp bins, which have approximately the size of a nucleosome plus spacer (Li and Reinberg, 2011). The average counts of reads mapping in each 200-bp bin in the genome (bigWig format) were used as the input for S3V2-IDEAS (Kent *et al*., 2010; Ramírez *et al*., 2016). The package accepts as input any bigWig files that can be represented as mean of mapped reads per bin. For example, starting with a set of bigWig files from BLUEPRINT data portal (Martens and Stunnenberg, 2013) or a set of bigWig converted from bam files by deeptools (Ramírez *et al*., 2016).

### 3. The S3V2 method for data normalization and denoising

The aim of the data normalization is to reduce the technical noise so that it cannot obscure the real biological differences. The recently developed S3norm method uses a non-linear transformation to match both the mean signals in the common peak regions and the non-zero mean signals in the common background regions between two data sets (Xiang, Keller, Giardine, *et al*., 2020). This new method can outperform other common normalization methods in tasks such as explaining gene expression levels or in consistency of peak calling between replicates (Xiang, Keller, Giardine, *et al*., 2020). In this package, we improved the S3norm method to improve the denoising of the datasets while normalizing them.

The key step of S3norm normalization is to match the mean signals in both the common peak regions and common background regions. However, for some data sets, we observed that matching only the mean signal is not sufficient to match the noise of the common background regions. As shown in Supplementary Figure 1A, the background noise of CTCF ChIP-seq in MONO cell type has a high standard deviation. As a result, even after matching the common background mean, the background noise in that data set is still much higher than the reference data sets (Supplementary Figure 1B and D). An extension of S3norm to simultaneously match the mean and standard deviation of the background regions would be expected to reduce the background noise to a similar level across different data sets (Supplementary Figure 1C and D).

The aim of S3norm is to produce data normalized across an entire genome, and thus it must be able to simultaneously incorporate and process the signal in both peak regions and background regions. A straightforward way to reduce the background noise without influencing the peak regions is to first split the genome into peak regions and background regions, and then normalize each of the two kinds of regions separately. However, this simple method can potentially create signal gaps between the two kinds of regions which do not reflect the original data. To overcome this signal gap issue, we incorporated a concept from the previous methods, in which the mapped reads in the same genomic region can be split and analyzed as the reads coming from different epigenetic structures, such as the foreground and background (Mahony *et al*., 2014) or the nucleosome free regions and accessible nucleosomes (Tarbell and Liu, 2019). In the improved version of S3norm (S3V2), we built on this idea by first splitting the reads in **each 200-bp bin** into the reads inferred to come from the foreground component and the reads inferred to come from the background component. Then, we used the two models (foreground and background) to normalize the two components separately, so that both the non-zero means and non-zero standard deviations for the background components can be matched.

Let *RC_i,ref_* and *RC_i,tar_* be the observed average read counts in bin *i* in a reference data set and a target data set. Similar to the quantile normalization (Bolstad *et al*., 2003), we used the average read counts of all data sets as the reference data set for each epigenetic feature. The reference data sets of different epigenetic features were first normalized to a same signal scale by the original S3norm method which can match the mean signals in the foreground regions and the non-zero mean signals in the background regions by a nonlinear transformation model (Xiang, Keller, Giardine, *et al*., 2020).

Then, for each epigenetic feature, we split the signals in both the reference data set and all individual data sets (target data set) into two components. To reduce the computational complexity, we used a relatively conservative but simple and fast approach in this step. Specifically, we estimated the number of reads coming from the background component by using the highest average read count in the non-peak region within 1kb around each 200-bp bin.

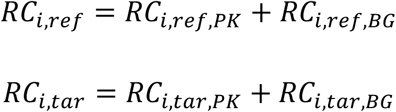

where *RC_i,ref,PK_* and *RC_i,tar,PK_* denote the reads coming from the foreground component, and *RC_i,ref,BG_* and *RC_i,tar,BG_* denote the reads coming from the background component. When there is a weak false negative peak at the local regions, this method may overestimate the reads in the background component. But on the other hand, it is less likely to inflate the background noise which can prevent identifying problematic epigenetic states with epigenetic features that contradict with each other in the downstream data integration module.

We then applied an exponential regression model to match the *RC_i,ref,PK_* and *RC_i,tar,PK_* between the two data sets.

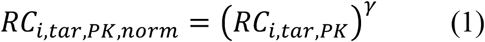

where the *γ* is the parameter learned from the regression model for the foreground component normalization. Here, we used the exponential regression model because it can guarantee there is no signal gap created between the peak signal and background signal after the normalization.

For the *RC_i,ref,BG_* and *RC_i,tar,BG_*, we used the following model to match the non-zero means and non-zero standard deviations between the two data sets.

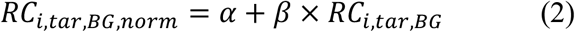

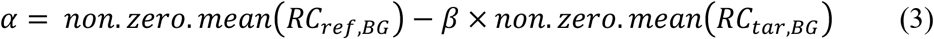

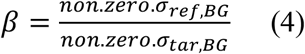

where the *α* and *β* are the two parameters for the background component normalization and *σ* is the standard deviation of the non-zero values in the backgrounds. Since the data sets with higher background noise usually have higher non-zero standard deviations, matching the nonzero standard deviations can greatly denoise these data sets. As a result, the overall background signal can be reduced to a similar level across all data sets (Supplementary Fig. 2A versus B). One expectation of a substantial reduction in background across the datasets is that the consistency in signal (including the background) between replicates for an epigenetic feature would be increased. This expectation was confirmed by the improvement of the consistency of epigenetic signals between two biological replicates that is measured by R-squared (Supplementary Fig. 2C).

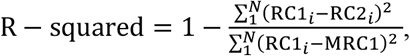

where the RC1_*i*_ and RC2_*i*_ are the S3V2 normalized average read counts in bin *i*, MRC1 is the whole genome mean of all RCl_*i*_, and N is the number of 200-bp bins.

This improvement in consistency was also observed when comparing the epigenetic states for each replicate after genome segmentation using IDEASs (Supplementary Fig. 2D). After normalizing each component (foreground and background) separately, we generated the final S3V2 signal tracks by adding those two components together in each 200 bp window.

In the S3V2 method, we trained the models by only using information in the common peak regions and common background regions, thereby learning values for the parameters *γ, α*, and *β* in equations (1)-(4). This is based on two assumptions made in previous studies (Shao *et al*., 2012; Xiang, Keller, Giardine, *et al*., 2020). First, we assumed the common peaks tend to regulate processes occurring in all cell types, such as the expression of constitutively active genes, so that their mean signal be similar after normalization. Secondly, the signals in common background regions are technical noise which should also be equalized after normalization. Then we applied those trained models to all regions along the whole genome.

In ChIP-seq analysis, the dynamic Negative Binomial (NB) distribution is often used to model the local background in each data set. Here, we also adjusted the local background in these S3V2 normalized signals by transforming them to −log10 p-value based on a dynamic background model with NB distribution (Xiang, Keller, Giardine, *et al*., 2020; Zhang *et al*., 2008).

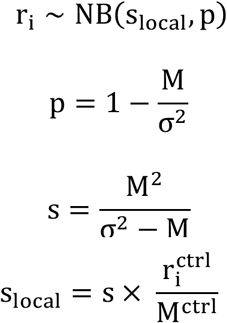

where p denotes the probability of success parameter, and s_local_ denotes a shape parameter of the in the dynamic NB model, the 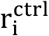 denotes no antibody control signals at 200-bp bin i, and the M^ctrl^ denotes global mean of no antibody control signals. For each bin i, the 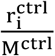 is used to capture the variation of local background. Here, the non-zero means and non-zero variances of different no antibody control signals are first scaled to the same level. Since S3V2 can match both the non-zero mean and non-zero variance of the background regions, the baseline parameters (M and *σ*^2^) in the model are estimated by only using the reference data set. Such a design can guarantee that the relative difference in the normalized average read counts are not changed in the −log10 p-value transformation.

### 4. Identifying epigenetic state and signal intensity state by IDEAS

Genome segmentation methods have been widely used to integrate and interpret the multidimensional epigenomic data sets in different cell types (Ernst and Kellis, 2012; Hoffman *et al*., 2012; Zhang *et al*., 2016; Xiang, Keller, Heuston, *et al*., 2020; Zhang and Hardison, 2017). Previous studies have demonstrated a 2-D (along the genome and across cell types) IDEAS genome segmentation model can perform better than other genome segmentation methods in both the consistency of state assignment and the ability to handle missing information (Zhang *et al*., 2016; Zhang and Mahony, 2019). In this package, we incorporated the IDEAS model as the second module to integrate the multi-dimensional epigenomic data sets into state tracks which are more interpretable and applicable.

The states learned in the IDEAS system can reflect multiple or single features, and we introduced two modes in this module to handle this versatility. The first mode is called epigenetic state mode (ES mode). When there are multiple epigenetic features in the input, the package can use the IDEAS model to integrate those multi-dimensional signals into a set of distinct epigenetic states (Supplementary Fig. 3). In this mode, each epigenetic state is represented by a unique color that is created by mixing the colors representing each individual epigenetic feature.

Moreover, since the IDEAS model is designed in a Bayesian framework, the state models can be easily incorporated as prior information in new IDEAS data integration analysis (Supplementary Fig. 3). In S3V2-IDEAS GitHub, we saved several epigenetic state models with different combinations of epigenetic features, so that user can easily use them as prior information to integrate their own epigenomic data sets.

The second mode is called signal intensity state mode. When only one epigenetic feature is covered in the input data, the package can use the IDEAS model to discover the commonly occurring signal intensity states (Supplementary Fig. 4A and B), which can provide detailed information about the epigenetic modification. To facilitate comparisons across different studies, we fixed the number of intensity states to 4. As shown in Supplementary Figure. 4C, the signal intensity states can be used to distinguish the peak region from the peak shoulders which can facilitate the downstream extraction of master peak list across multiple cell types. In this mode, we used 50-bp as the bin size because it can facilitate downstream identification of narrow peaks.

After identifying the states, both modes can generate a genome segmentation track by assigning a state to each genomic location in each cell type based on the signal pattern. In the IS mode, we also generated an additional signal intensity state track based on the average normalized signal track. This additional average signal intensity state track can be used to extract master peak list in the downstream analysis (see next section).

In our previous analysis, we noticed that the IDEAS model can generate an epigenetic state by merging some rare patterns as one epigenetic state, which we referred as heterogeneous states (Xiang, Keller, Heuston, *et al*., 2020). As a result, these heterogeneous states often show the combinations of epigenetic modifications that are unlikely to exist, e.g. by merging multiple, rarely-observed combinations. To avoid this confounding information, we followed our previous work (Xiang, Keller, Heuston, *et al*., 2020) and incorporated into the package a script to check the coefficient of variation 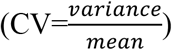 of each epigenetic state. For the states with higher CV, we suggest that the user to remove them and then use the remaining states as prior information to rerun IDEAS. In this process, those genomic regions with rare patterns can be assigned to a most similar epigenetic states that occurring frequently.

### 5. Extracting master peak list across multiple cell types

A consensus master peak list is often used to analyze the epigenetic modifications across different cell types (Xiang, Keller, Heuston, *et al*., 2020; Lun and Smyth, 2016; Meuleman *et al*., 2019). Although simply pooling and merging the peaks in multiple cell types has been widely used, the merging step may create broader peaks that do not accurately reflect the positions of the epigenetic modifications, especially when one or a few data sets have very strong signals. In the example shown in Supplementary Figure 5, based on the normalized ATAC-seq signal, there should be one candidate cis-regulatory element (cCRE) (between the gray dashed lines) at the promoter of *Hba-a1* gene. However, the master peak generated by pooling and merging the MACS2 peaks in all cell types covers the entire *Hba-a1* gene region, thereby obscuring the separate cCRE at the promoter regions (between the blue dash lines).

To address this issue, several strategies have been developed to extract master peak lists (Meuleman *et al*., 2019; Moore *et al*., 2020). In the S3V2-IDEAS, we introduce a novel hierarchical method to systematically extract a master peak list that preserves the resolution to pinpoint positions of the epigenetic modifications in each cell type.

Since merging the peaks in different cell types tends to broaden the inferred peak calls, we avoided merging while extracting the master peaks, instead taking advantage of the signal intensity state tracks. Specifically, we first generated an average signal track and the corresponding signal intensity track by using the S3V2 normalized signals of all cell types (Supplementary Fig. 6B). Here, this simple averaging step can first reduce the noise in each cell type. We then collected the genomic regions that are assigned with the strongest signal intensity states into the list of master peaks (MPs) (Supplementary Fig. 6A-C). This initial list derived from the analysis of the signal intensity states in the average signal track will reveal primarily the peaks that are present in many cell types. The average signal track dilutes the strong peak signals that are present in only one or a few cell types, so to avoid missing the cell type specific peaks, we also collect the genomic regions with the strongest signal intensity states in each cell type that do not intersect with peaks already in the MP list (Supplementary Fig. 6A and C). This hierarchical design can guarantee the peaks identified based on the average signal track have the highest priority to be selected as the master peaks. To get the final list of MPs, we repeat this collection process for each weaker signal intensity states so that the relatively weaker MPs can also be collected.

Comparing with the MP list identified by pooling and merging the MACS2 peaks in all cell types, the MPs identified by our package are significant narrower (Wilcox -test p-value < 2.2e-16), so that they can pinpoint the position of the epigenetic modifications across different cell types with higher accuracy (Supplementary Fig. 7).

## Supplementary Figures

**Supplementary Figure 1.**
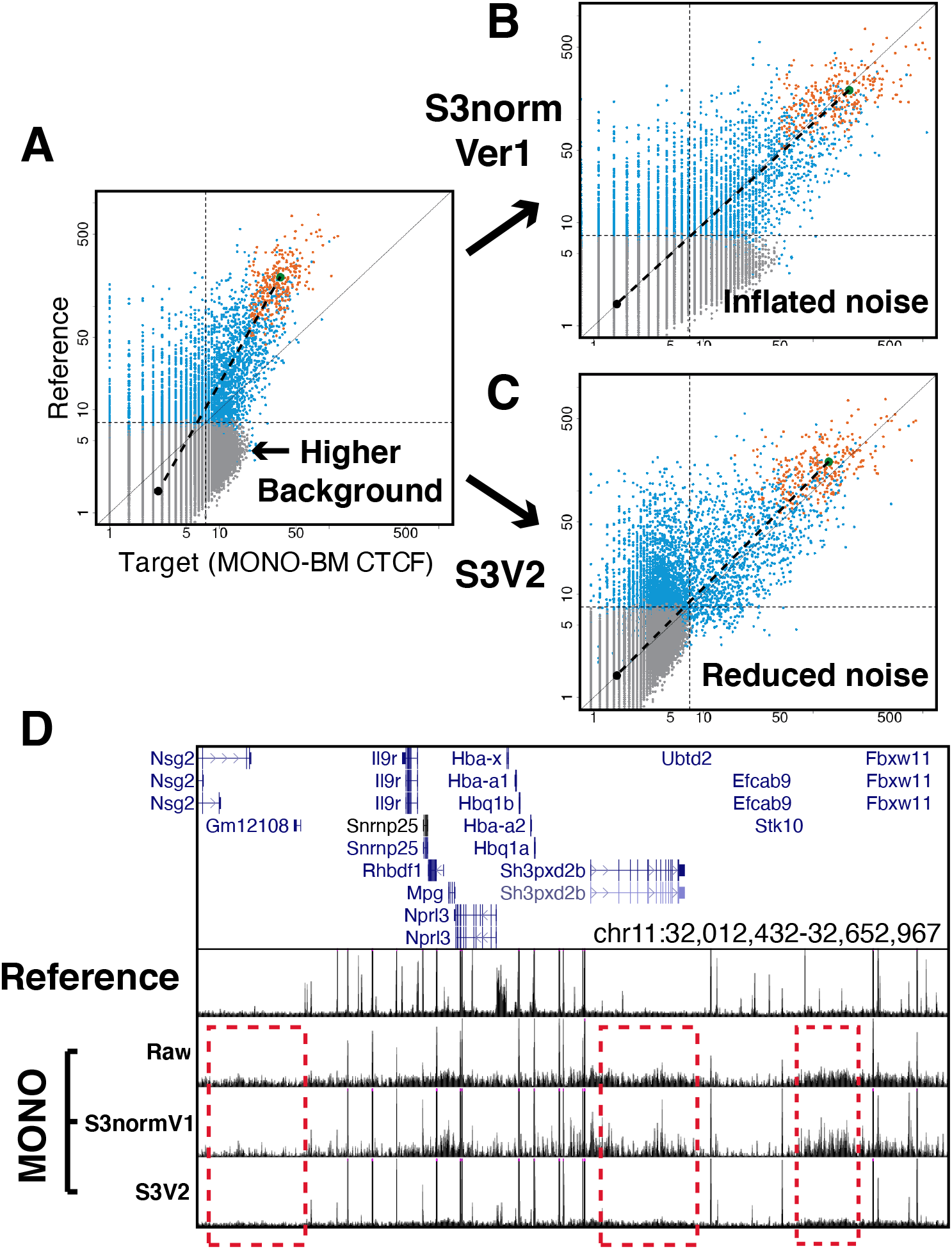
The first version of S3norm does not equalize the background signal in data sets with higher background variance. The scatterplots of the average read counts before (A) and after (B) S3norm between the Reference data set and the monocyte in bone marrow (MONO-BM) data set. (C) On the contrary, the S3V2 can successfully reduce the noise of the same data set to the level of the Reference data set. (D) The corresponding signal tracks of the Reference data set and the MONO-BM data set. The red dashed boxes enclose regions with inflated noises after first version of S3norm, but reduced noise after S3V2.

**Supplementary Figure 2.**
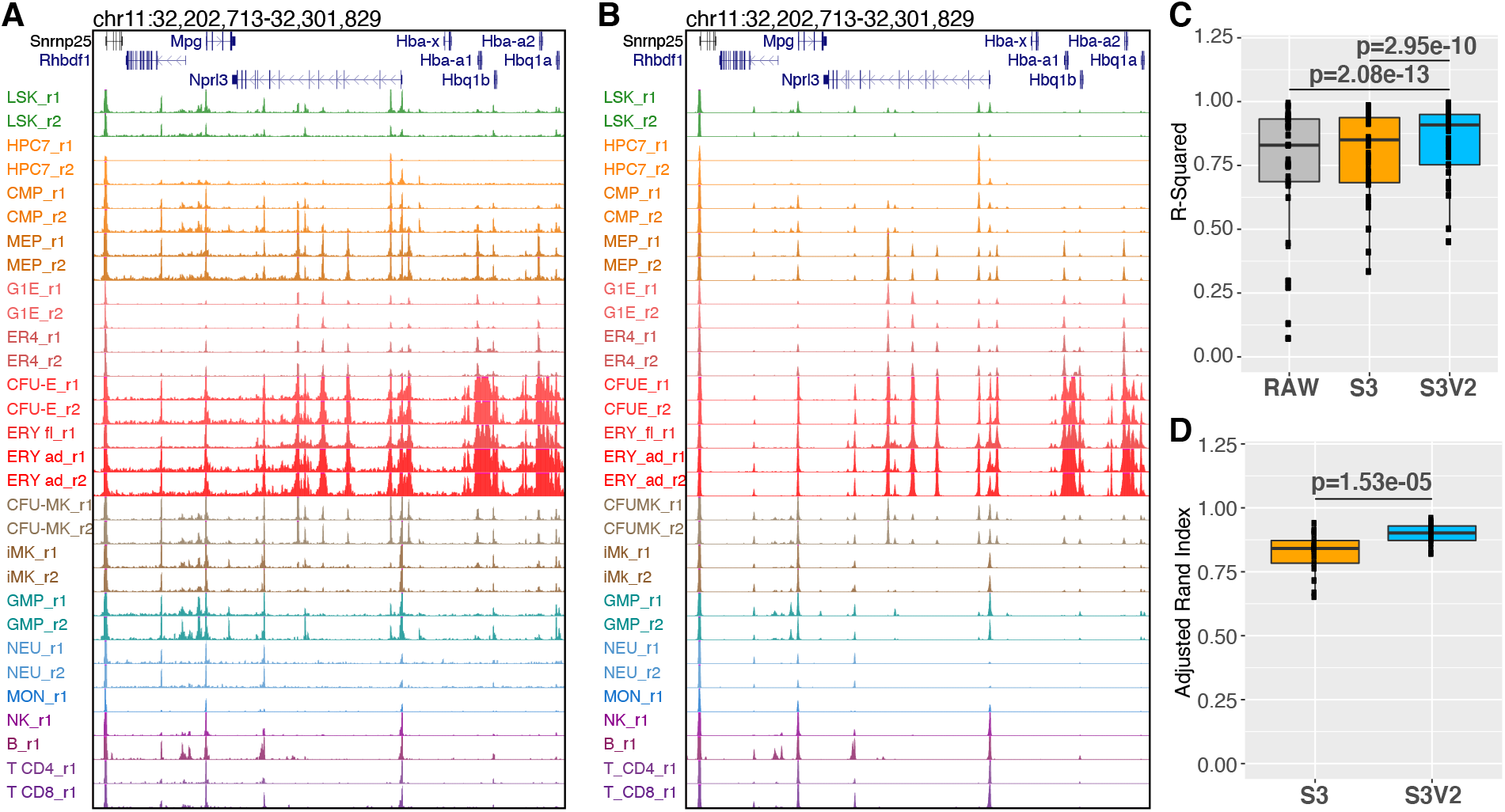
The ATAC-seq signal before (A) and after S3V2 normalization (B). The different colors represent different cell types. The two biological replicates were label as r1 and r2. (C) The R-squared is used to measure the consistency of epigenomic signals between biological replicates. (D) The Adjusted Rand Index is used to measure the consistency of epigenetic states between biological replicates.

**Supplementary Figure 3.**
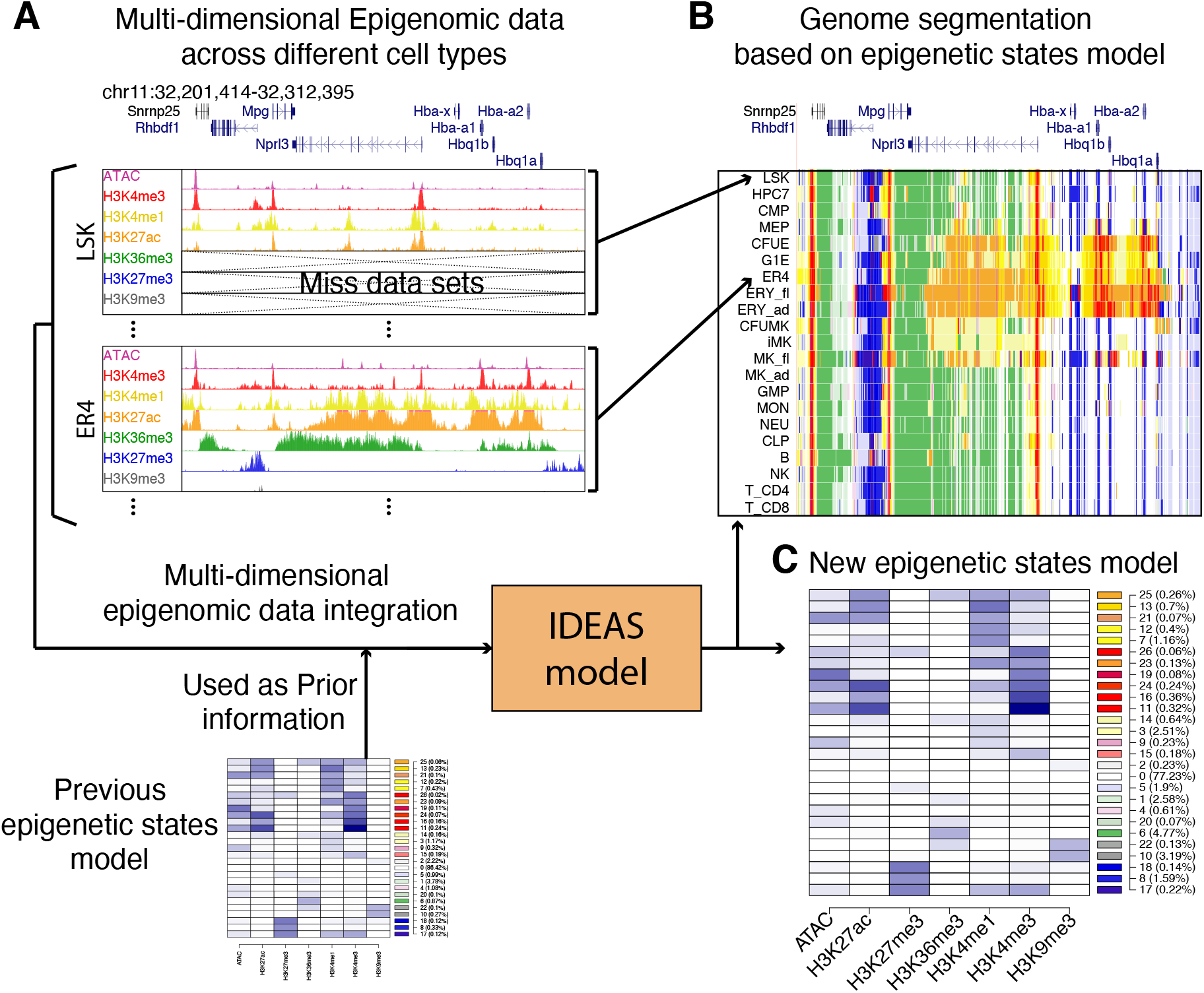
The ES mode can integrate multiple epigenetic features in multiple cell types as epigenetic states model. (A) The S3V2 normalized epigenomic signals in different cell types. The missing information in certain cell types can be handled in IDEAS model by borrowing information from similar cell types. (B) and (C) The IDEAS model can integrate the epigenomic signals as epigenetic states model and the corresponding genome segmentation tracks. The epigenetic states model generated in previous studies can be easily incorporated as prior information in the IDEAS model.

**Supplementary Figure 4.**
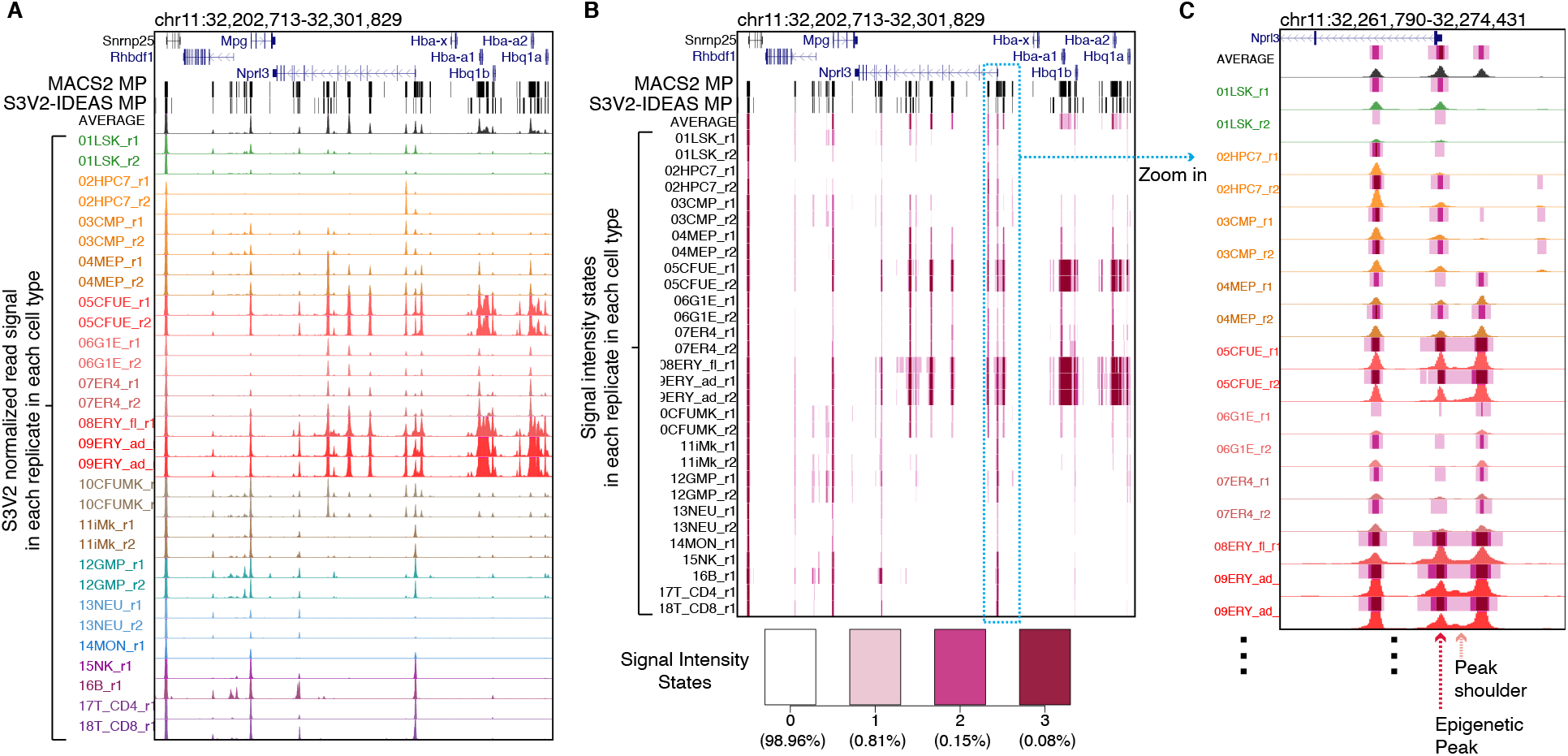
The IS mode can integrate the signals of the same epigenetic feature as signal intensity state tracks. (A) The S3V2 normalized ATAC-seq signal in each replicate of each cell type. (B) The signal intensity state tracks based on the normalized ATAC-seq signals. The Master Peak (MP) lists generated by MACS2 peak calling method and the S3V2-IDEAS pipeline are show on top of each figure. (C) The signal intensity state and the corresponding S3V2 normalized ATAC-seq signals.

**Supplementary Figure 5.**
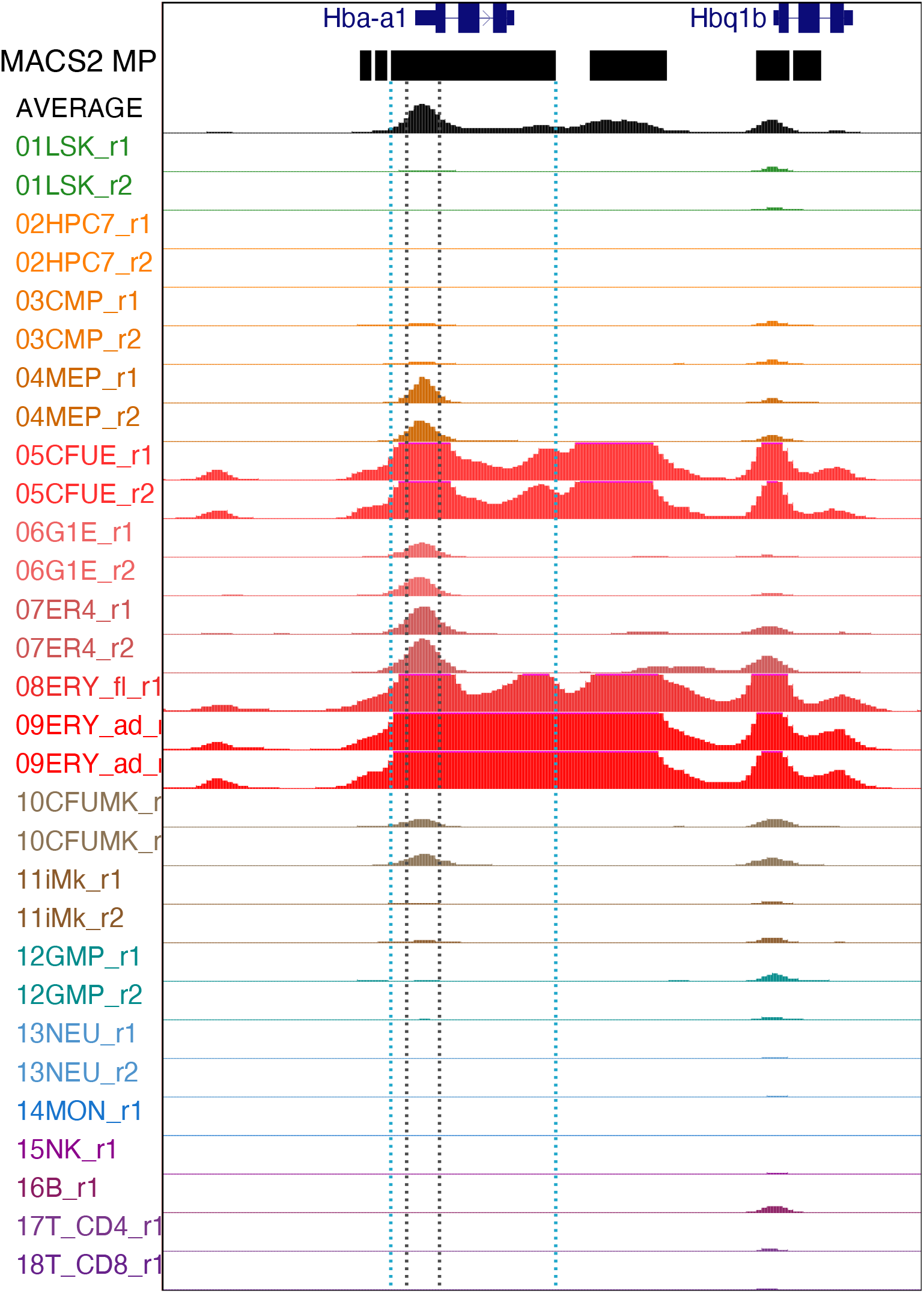
The master peaks generated by pooling and merge MACS2 peaks in all cell types. The region between the dark gray dash lines is the accessible region at the promoter of *Hba-a1* gene across multiple cell types. The region between the blue dash lines is the master peak identified by pooling and merging the MACS2 peaks in all cell types.

**Supplementary Figure 6.**
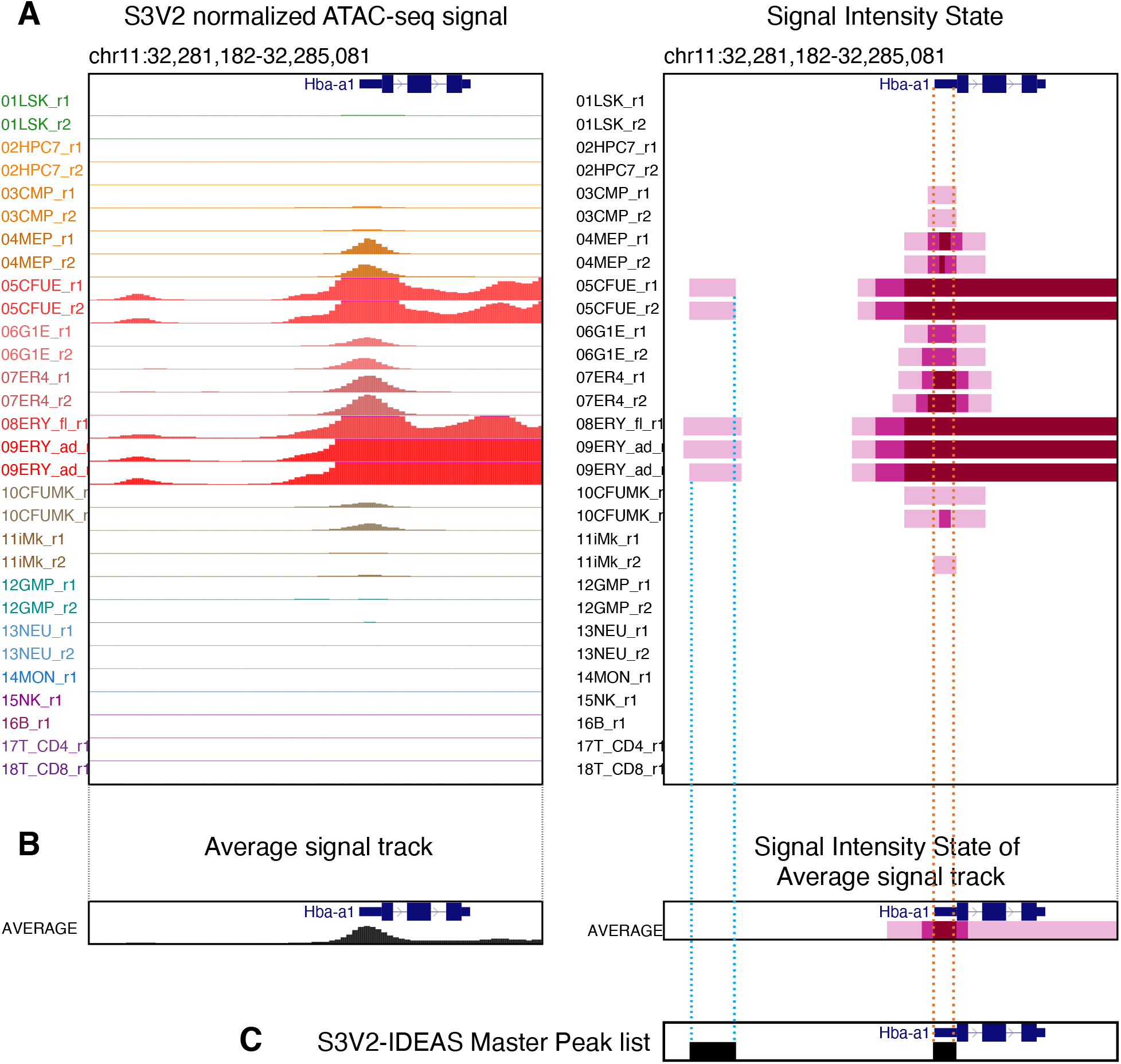
The hierarchical method for extracting master peaks from signal intensity state tracks. (A) The S3V2 normalized signals and the corresponding signal intensity state tracks in multiple cell types at the *Hba-a1* gene locus. (B) The average signals and the corresponding signal intensity state tracks generated using the information in all cell types. (C) The peaks are extracted first from the average signal intensity state track (the peak between the orange dash line) and then from the peaks extracted from individual cell type (the peak between the blue dash line) that do not intersect with the peaks of the average signal intensity state track. These sets of peaks are combined to generate the master peak list in S3V2-IDEAS pipeline.

**Supplementary Figure 7.**
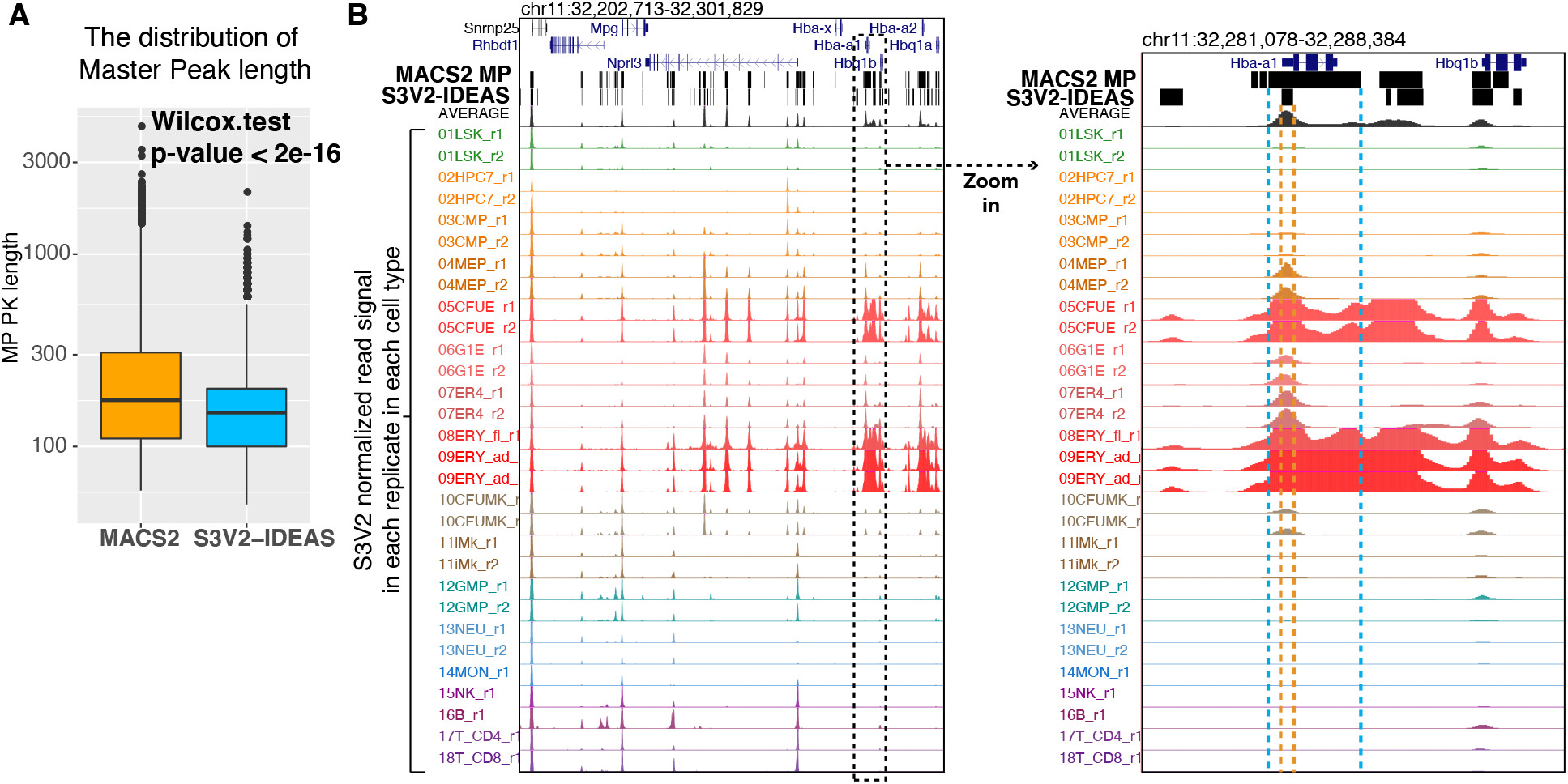
Comparing the master peak lists generated by S3V2-IDEAS and MACS2. (A) The distributions of peak lengths of master peaks generated by S3V2-IDEAS pipeline and MACS2 are shown as box-plots. (B) The master peak at the *Hba-a1* gene locus. The region between the orange dash lines is the master peak identified by S3V2-pipeline. The region between the blue dash lines is the master peak identified by pooling and merging the MACS2 peaks in all cell types.

## Notes

### Competing Interest Statement

The authors have declared no competing interest.

